# Global Sesame Germplasm Harbors Untapped Genetic Potential for Crop Improvement

**DOI:** 10.64898/2026.09.02.748780

**Authors:** Sookyeong Lee, Sae Hyun Lee, Ju-Young Ahn, Jae-Eun Lee, SuKyeung Lee, Murukarthick Jayakodi, Gi-An Lee, Tae-Jin Yang

**Affiliations:** National Agrobiodiversity Center, National Institute of Agricultural Science, Rural Development Administration, Jeonju 54874, Korea; Department of Agriculture, Forestry and Bioresources, Plant Genomics and Breeding Institute, Research Institute of Agriculture and Life Science, College of Agriculture and Life Sciences, Seoul National University, Seoul 08826, Korea; Department of Applied Plant Sciences, Division of Bioresource Sciences, Kangwon National University, Chuncheon, 24341, Republic of Korea; Texas A&M AgriLife Research and Extension Center, Dallas, TX, USA

**Author notes:** Corresponding author: Murukarthick Jayakodi, Gi-An Lee, and Tae-Jin Yang. These authors contributed equally to this work (Sookyeong Lee, Sae Hyun Lee).

**Keywords:** sesame, germplasm, flowering, lignan, antioxidant, GWAS, *k*-mer, diversity, Genebank

## Abstract

**Background:** Sesame (*Sesamum indicum* L.) is one of the oldest oilseed crops. Its seeds accumulate lignans and antioxidants that determine their nutritional value. Global demand for sesame is rising rapidly, but climate change increasingly threatens sesame yields and seed quality. Here, we analyzed a worldwide panel of 300 sesame accessions to explore the genetic basis of key adaptive and quality traits, specifically flowering time, seed lignan content, and seed antioxidant capacity.

**Results:** Whole-genome resequencing revealed previously unreported genetic diversity, expanding the resources available for sesame breeding. By integrating *k*-mer-based genome-wide association analysis with a graph-based sesame genome, we identified structural variants associated with differences in flowering time and lignan accumulation. A 9.4-kb deletion on chromosome 6 that disrupted SIN_1018434 (an ortholog of Arabidopsis *PHOTOPERIOD-INDEPENDENT EARLY FLOWERING 1*) and a 6.2-kb *Copia*-type retrotransposon insertion upstream of SIN_1004470 on chromosome 11 were associated with early flowering. The intact alleles at both loci were associated with delayed flowering and were predominant in low-latitude accessions. High seed lignan content was associated with non-synonymous mutations and copy number variants in glycosyl hydrolase genes on chromosome 6 and an 8.2-kb deletion spanning SIN_1019378 on chromosome 13. Association signals for seed antioxidant traits coincided with loci involved in abiotic stress responses and seed-coat pigmentation.

**Conclusions:** These findings highlight the importance of exploring diverse germplasm to uncover previously unrecognized adaptive and quality-associated genomic variation. The loci identified here provide molecular targets for developing climate-adapted cultivars with improved seed quality.

## Background

Sesame (*Sesamum indicum* L.), one of the earliest oilseed crops[1–3], was likely domesticated from the wild progenitor Malabar sesame (*S. indicum* subsp. *malabaricum*)[3–5]. Modern sesame belongs to the family Pedaliaceae and is a predominantly self-pollinating diploid species (2n = 2x = 26)[6, 7]. Although the genus *Sesamum* currently comprises 31 accepted species[8], *Sesamum indicum* is the only one widely cultivated as an oilseed crop[4, 9, 10]. Archaeobotanical evidence indicates that sesame was already being cultivated in northwestern South Asia by the Mature Harappan period of the Indus civilization, approximately 2500–2000 BC[1, 3]. This long history of cultivation reflects its agronomic utility as well as its nutritional and industrial value to human societies.

Sesame seeds are rich in oil (∼58% of dry seed weight), protein (∼25%), and antioxidants; sesame oil is valued in food processing because of its high oxidative stability and distinctive lignan composition[4, 11–13]. Lignans are a class of phenylpropanoid-derived plant specialized metabolites[14, 15], and the major lignans in sesame, sesamin and sesamolin, are benzodioxol-substituted furofurans reported to exhibit various biological activities, including antioxidant and anticancer effects[11, 16]. Driven by the increasing demand for sesame products, the global harvested area expanded from 4.96 million ha in 1961 to 12.97 million ha in 2021, while total production increased from 1.42 million tons to 6.67 million tons[12, 17–19]. However, despite these substantial expansions in harvested area and total production, seed yield per unit area has not improved substantially[18, 20], and climate change continues to threaten the stability of both yield and quality[21, 22]. In particular, sesame yield and lignan content are strongly influenced by abiotic stress[16, 21–24]. Adjusting to the more extreme environmental conditions brought about by climate change requires breeding strategies that maintain stable yield and quality across diverse and fluctuating seasonal conditions.

Advances in genomics have transformed crop improvement by enabling the dissection of complex agronomically important traits. Declining sequencing costs facilitate genotyping and trait mapping in crop species including sesame, allowing the identification of genetic determinants underlying yield, oil-related traits, and responses to environmental stresses[25–33]. Recent studies have highlighted the importance of structural variation in addition to simple single-nucleotide polymorphisms (SNPs) in the regulation of agronomic traits, prompting the development of pangenome frameworks[34–39]. In sesame, an intraspecific pangenome study[40], an intraspecific multi-reference genome study[41], and a genus-level pangenome study[9] collectively analyzed genome assemblies representing nine distinct cultivated *S. indicum* accessions and six wild *Sesamum* species. The latter two studies identified structural variants associated with plant architecture, yield-related traits, and resistance to Fusarium wilt[9, 41]. Importantly, diverse and representative accessions integrated with genomic approaches are essential for agronomic improvement and climate adaptability. Previous reports characterized the variation in days to flowering in a global panel of natural sesame accessions, revealing variation ranging from fewer than 30 days to over 90 days[17, 25, 26]; this diversity may have facilitated the expansion of sesame cultivation across latitudes[17, 26, 42]. In addition, natural variation in lignan content and antioxidant capacity among sesame accessions suggests that they represent a vast genetic resource for improving seed quality and stability through breeding[43–47]. We reasoned that this diversity provides a vital opportunity to elucidate the genetic basis of climate adaptability and quality-related traits in sesame.

In this study, we quantified flowering-related traits that contribute to cultivation flexibility, as well as seed-quality traits for the stable production of high-quality oil, including contents of the lignans sesamin and sesamolin, total phenolic content, and antioxidant activity, in 300 sesame accessions collected worldwide. Using resequencing data from these accessions, we performed a *k*-mer-based genome-wide association study (GWAS) and identified genetic variants associated with flowering and seed quality using a graph-based genome. The variants identified here can be directly leveraged to develop marker-assisted selection strategies for key agronomic and quality traits, including flowering time and seed lignan content, and provide a genetic foundation for future studies and breeding aimed at regulating flowering and lignan metabolism in sesame.

## Results

### Population structure of global sesame accessions reflects the dynamic trading history of sesame

To characterize genome-wide variation in sesame, we selected 300 globally representative accessions fom the RDA-Genebank and subjected them to whole-genome resequencing at approximately 20-fold genome coverage (Additional file 1: Fig. S1; Additional file 2: Tables S1 and S2). After mapping reads to the Zhongzhi No. 13 reference genome (v2)[48], we identified ∼7.8 million (M) variants, consisting of SNPs and insertions/deletions (InDels) across the genomes of these 300 accessions.

Using 1.9 M high-quality SNPs with a minor allele frequency above 5% and a missing genotype rate below 20% (Additional file 1: Fig. S2), we performed population structure and phylogenetic analyses (Fig. 1a, b). At K = 2, the population structure analysis identified two major ancestry components (Fig. 1a). Based on the relative proportions of these ancestry components, the accessions were classified into two predominant ancestry groups, designated G1 and G2, and an admixed group, designated G3. G1, G2, and G3 consisted mainly of accessions from Eurasia, Africa, and the Americas, respectively. Notably, the major ancestry components did not follow a simple binary pattern of Asian and African origin, suggesting that the present-day global sesame gene pool has been shaped by geographical origin as well as post-domestication dispersal and admixture.

**Figure 1.**
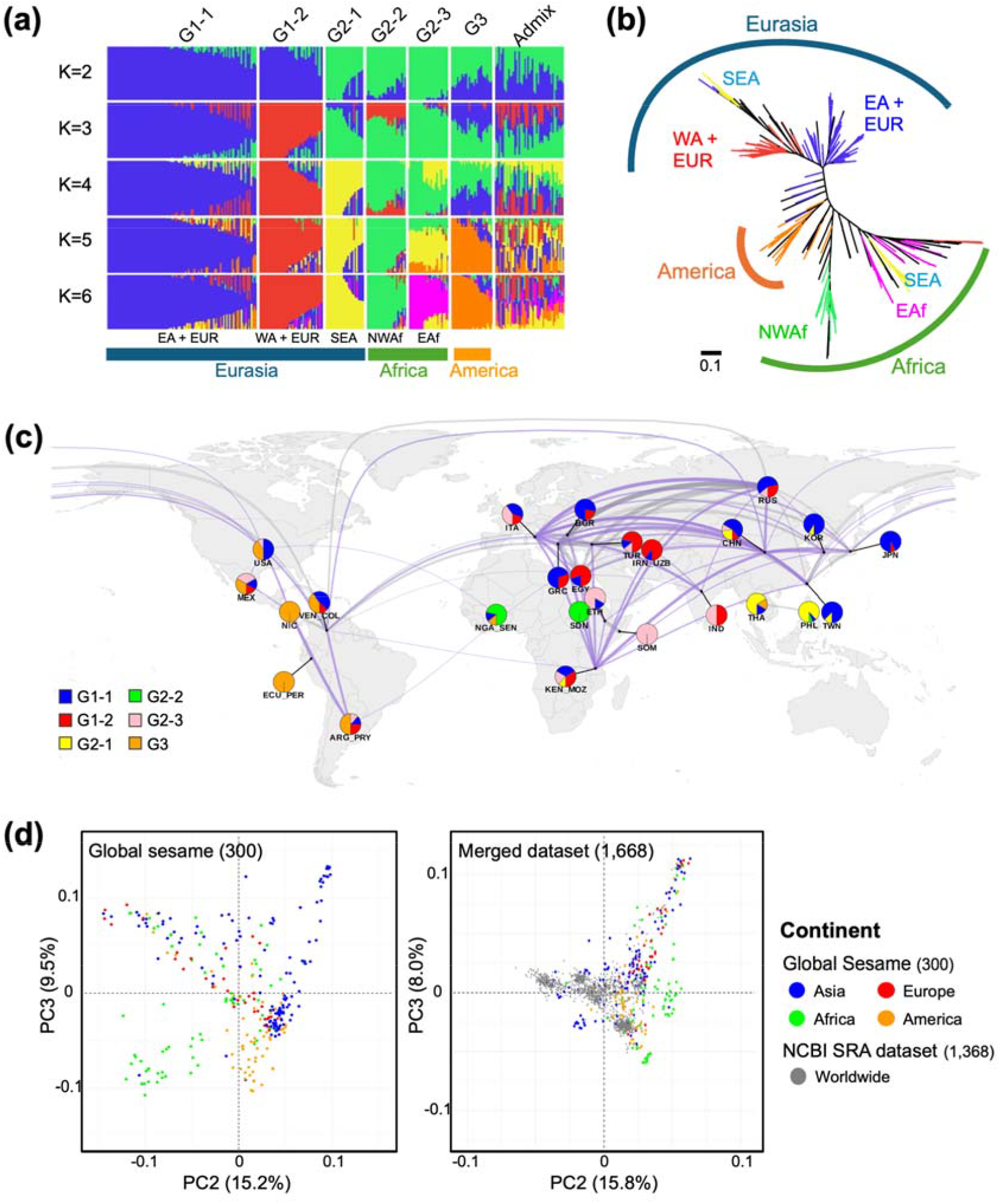
Population structure, phylogenetic relationships, and principal component analyses of 300 sesame accessions. (a) Population structure analysis of the 300 sesame accessions using ADMIXTURE at *K* = 2–6. Each vertical bar represents one accession, and colors indicate inferred ancestry components. Accessions are grouped by continent of origin. (b) Phylogenetic tree of the 300 sesame accessions. Branch colors indicate the six subgroups identified at *K* = 6, with black branches indicating admixed accessions. (c) Regional genetic overlap and population structure of global sesame accessions. Pie charts show the *K* = 6 ADMIXTURE ancestry of the 25 collection regions. Within each chart, slice size is proportional to the number of accessions assigned to a population group (G1-1, blue; G1-2, red; G2-1, yellow; G2-2, green; G2-3, pink; G3, orange). Charts represent the 253 confidently assigned accessions, excluding admixed accessions. Curved lines connect regions with high regional genetic overlap (RGO **≥** 0.7, ReMIXTURE run 10), with thicker lines indicating stronger overlap. Purple lines indicate that at least one of the two connected regions is high diversity, defined as the most diverse 30% of regions, and grey lines indicate the regions are not high diversity. (d) Principal component analysis (PCA) of sesame accessions based on PC2 and PC3. Left, PCA of the 300 global sesame accessions. Right, PCA of the combined dataset comprising the 300 global accessions and 1,368 publicly available accessions from the NCBI Sequence Read Archive (SRA). EA; East Asia, EUR; Europe, WA; West Asia, SEA; Southeast Asia, NWAf; North and West Africa, EAf; East Africa, Admix; Admixture.

Inspection of ancestry patterns across increasing *K*-values further highlighted this complexity. At *K* = 6, the broad ancestry classes observed at *K* = 2 were resolved into six genetic subgroups. Each subgroup was assigned to one of the broad classes, resulting in two G1-related subgroups (G1-1 and G1-2), three G2-related subgroups (G2-1, G2-2, and G2-3), and one predominantly admixed subgroup (G3). The six genetic subgroups also broadly corresponded to the geographical origins of the accessions. The G1-related subgroups comprised most Asian accessions, excluding many from Southeast Asia, along with European accessions (Fig. 1a). The persistent clustering of European and Asian accessions together suggests substantial genetic proximity between these groups and is consistent with long-term exchange of sesame materials across Eurasia[1, 3]. Similarly, the G2-related subgroups included African accessions together with a substantial proportion of Southeast Asian accessions. This pattern suggests that the current genetic relationship between Southeast Asian and African accessions may reflect repeated movement of materials across connected regions.

We observed a more complex structure in Southeast Asian accessions. In the ADMIXTURE analysis, many Southeast Asian accessions showed greater similarity to African accessions than to other Asian accessions. This pattern was also supported by the phylogenetic tree, in which Southeast Asian accessions were distributed across both Eurasian-related and African-related lineages (Fig. 1a,b). Rather than forming a single regional lineage within the broader Asian gene pool, Southeast Asian accessions appear to retain multiple ancestral backgrounds, indicating that geographical origin alone does not fully predict genetic relatedness across sesame accessions. Environmental selection may also have contributed to these patterns. For example, agronomic traits such as days to flowering and days to maturity are strongly influenced by temperature and photoperiod[17, 49], raising the possibility that latitude-related selective pressures favored genotypes already suited to tropical and subtropical environments. Similarly, the admixed ancestry observed in American accessions is consistent with their introduction from multiple Old World sources, followed by recombination and regional adaptation after dissemination into the Americas[2, 3, 50].

Visualizing the *K* = 6 ancestry composition together with the regional genetic overlap (RGO) network provided an additional geographical perspective on these patterns (Fig. 1c). Among the confidently assigned accessions, we observed extensive high-overlap connectivity among accessions from Old World regions, whereas comparable connections among accessions from the Americas were relatively limited. This pattern is consistent with a longer and more interconnected history of shared ancestry and the exchange of materials across Africa, Asia, and Europe. Connections between the Old and New Worlds were not evenly distributed but were concentrated in a limited number of regions. Within the resulting network, China and Italy appeared as relatively well-connected nodes in East Asia and Europe, respectively, whereas Venezuela was well connected to other nodes in the Americas. These connectivity patterns suggest that introductions of sesame materials into the Americas may have occurred through a limited number of source or intermediary regions, followed by subsequent redistribution within the New World. However, because RGO reflects genetic similarity between regional populations and does not resolve the direction or timing of dispersal, these connections should be interpreted as plausible patterns of shared ancestry and historical exchange rather than as definitive migration routes. Together, these results suggest that the structure of the present-day global sesame population reflects a long history of material movement.

Principal component analysis (PCA) further showed that the global sesame panel captures broad genetic diversity (Fig. 1d, Additional file 1: Fig. S3). Asian accessions occupied the widest region of the PCA space and overlapped with European accessions, further supporting the close relationship between these two groups (Fig. 1d, left). Accessions from Africa and the Americas partially overlapped with other groups but also formed distinct clusters, indicating that their inclusion in our dataset substantially expanded the diversity present among the 300 accessions (Fig. 1d, left). When combined with datasets for the genomes of 1,368 sesame accessions previously published at NCBI, the global sesame panel occupied a broader genomic space than the NCBI dataset alone (Fig. 1d, right), indicating that the collection assembled in this study captures previously uncharacterized variation and expands the allelic landscape available for downstream genetic analysis.

This interpretation was further supported by measuring linkage disequilibrium (LD) decay. In the global panel of 300 accessions, LD declined to half of its maximum at 52.46 kb (Additional file 2: Table S3), which is shorter than the 88–99 kb previously reported in sesame[7, 25, 51]. Although direct comparisons among studies should be made cautiously because estimates can be influenced by sample composition, marker density, and analytical criteria, the relatively rapid LD decay observed here is consistent with the elevated genetic diversity in the panel of 300 accessions. This feature is also expected to improve mapping resolution by reducing the physical size of linkage blocks.

Taken together, these results show that the global sesame collection assembled in this study captures both geographically structured and admixed variation shaped by long-term dispersal, regional exchange, and environmental adaptation. The broad genomic space and relatively short LD decay provide a valuable foundation for association mapping, conservation of genetic resources, and breeding strategies aimed at broadening the adaptive and agronomic potential of cultivated sesame.

### Structural variants from low-latitude sesame accessions are associated with flowering time

To identify genomic regions associated with flowering time, we performed two GWAS: one using all SNPs, and the other using *k*-mers, with the global panel of 300 sesame accessions (Fig. 2). We tested for association with days to flowering (DTF), days to maturity (DTM), height to the first capsule (HTFC), and capsule zone length (CZL). DTF and DTM values were strongly correlated with the plant architecture traits CZL (negative correlation) and HTFC (positive correlation), suggesting that flowering time influences overall plant architecture (Fig. 2a, b, Additional file 1: Fig. S4a, and Additional file 2: Table S4). Across all four traits, SNP-based GWAS identified a significant association on chromosome 4, whereas *k*-mer-based GWAS detected additional candidate peaks on chromosomes 3, 6, 11, and 13, while also capturing the signal on chromosome 4 (Fig. 2c). The strongest association signal mapped to chromosome 4 (17,190,000–17,420,000 bp) within an LD block containing the previously reported candidate genes SIN_1013026 (*FLOWERING LOCUS T* [*FT*]*-like*) and SIN_1013025 (*HEADING DATE 3A*), previously mapped to the LG11 region[52, 53] (Additional file 1: Fig. S5a and Additional file 2: Table S5). The same LD block contained SIN_1013020 (*ABSCISIC ACID INSENSITIVE 5* [*ABI5*]), suggesting that abscisic acid (ABA) signaling and FT-related pathways may contribute to the regulation of flowering time in sesame (Additional file 1: Fig. S5a), although the strong LD structure across this region currently limits the resolution of causal variants.

**Figure 2.**
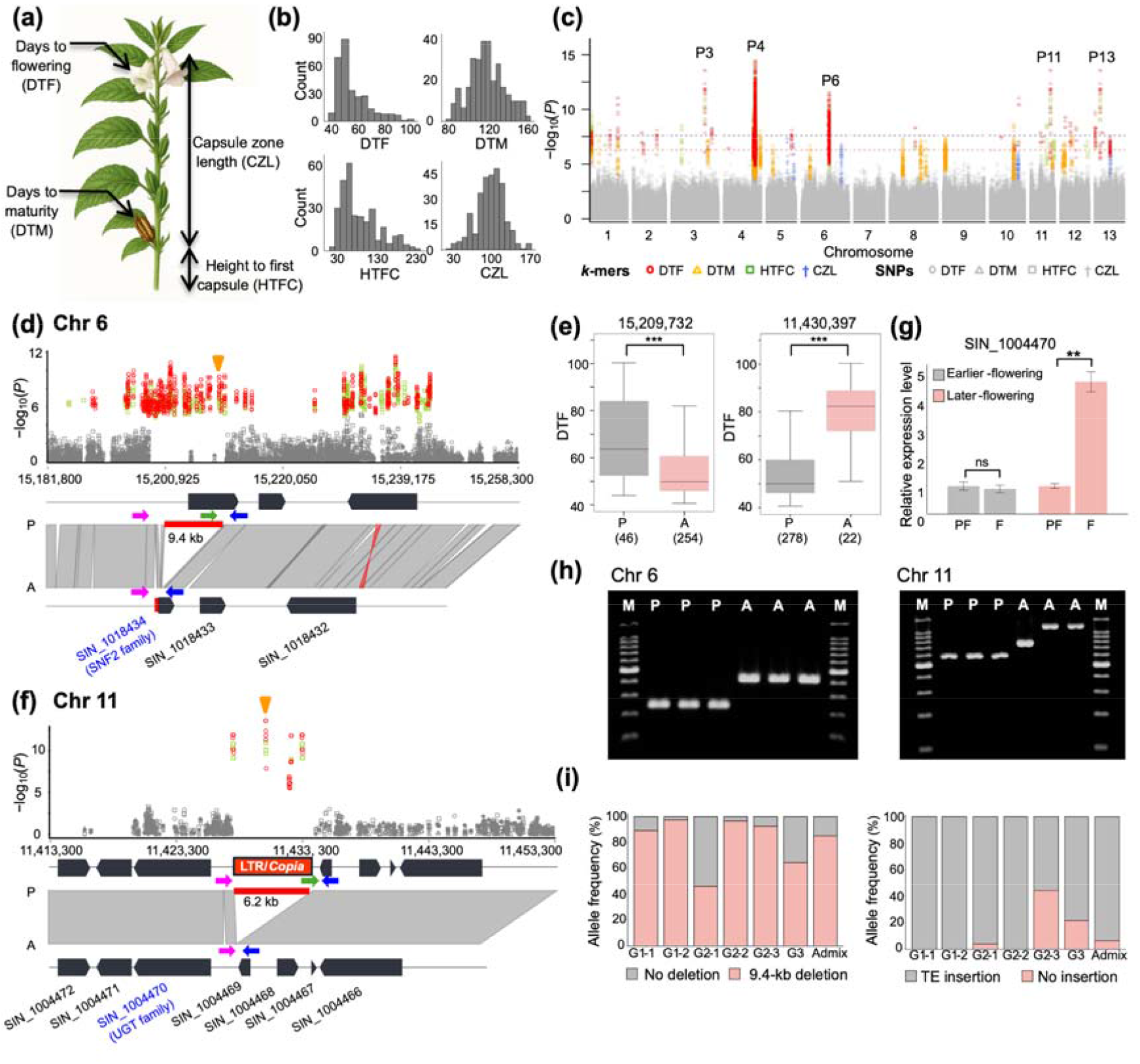
Phenotypic variation and identification of novel structural variants associated with flowering-related traits. (a) Schematic of days to flowering (DTF), days to maturity (DTM), height to first capsule (HTFC) and capsule zone length (CZL). (b) Distributions of DTF, DTM, HTFC, and CZL across three growing seasons. (c) Combined Manhattan plot of SNP-and *k*-mer-based GWAS using best linear unbiased estimates (BLUEs). Purple and pink dashed lines indicate Bonferroni thresholds for SNPs (*P* = 2.5 × 10**□□**) and *k*-mers (*P* = 5 × 10**□□**), respectively. (d) A 9.4-kb deletion on chromosome 6 includes the 5**′** region of SIN_1018434. The orange arrow at Chr6:15,209,732 bp indicates the start position of the significant *k*-mer. Primer arrows indicate non-deletion-allele-specific forward (green), common forward (pink), and common reverse (blue) primers. (e) Boxplots of DTF by presence/absence of the significant *k*-mer in the chromosome 6 SV (left) and chromosome 11 LTR/*Copia* region (right). Two-sided Wilcoxon rank-sum tests; \**P* < 0.05, \*\**P* < 0.01, \*\*\**P* < 0.001. (f) A 6.2-kb LTR/*Copia* retrotransposon on chromosome 11 is inserted between SIN_1004470 and SIN_1004469. The orange arrow at Chr11:11,430,397 bp indicates the start position of the significant *k*-mer. Primer arrows indicate TE–genome junction-specific forward (green), common forward (pink), and common reverse (blue) primers. (g) Relative expression of SIN_1004470 in leaves at pre-flowering (PF) and flowering (F) stages in early-flowering IT216140 (grey) and late-flowering IT201452 (pink). Means ± SE of three biological replicates. Student’s *t*-test; ns, not significant; \**P* < 0.05, \*\**P* < 0.01, \*\*\**P* < 0.001. (h) Gel electrophoresis of chromosome 6 (left) and 11 (right) SV markers in early- and late-flowering accessions. A 100-bp DNA ladder was used; expected amplicons are 240 bp (non-deletion) and 410 bp (deletion) on chromosome 6, and 584 bp (TE-insertion) and 785 bp (non-TE) on chromosome 11. (i) Distribution of chromosome 6 (left) and 11 (right) SVs among *K* = 6 subgroups defined in Figure 1a.

Notably, we detected a 9.4-kb deletion on chromosome 6 in the 5**′** region of SIN_1018434, an ortholog of Arabidopsis (*Arabidopsis thaliana*) *PHOTOPERIOD-INDEPENDENT EARLY FLOWERING 1* (*PIE1*, At3g12810), which is involved in *FLOWERING LOCUS C* (*FLC*)-mediated floral repression[24, 54]. This deletion encompasses 5.5 kb of genic sequence, resulting in a truncated gene and putative pseudogenization (Fig. 2d). Accessions carrying this deletion flowered significantly earlier than those without the deletion (Fig. 2e, left), consistent with the role of PIE1 as a floral repressor and the early-flowering phenotype observed in Arabidopsis *pie1* mutants[54].

Additionally, we detected significant identical *k*-mer peaks on chromosomes 3, 11, and 13, each mapping to an insertion of a *Copia*-type retrotransposon on the respective chromosome (Fig. 2c, 2f, and Additional file 1: Fig. S5b,c). Among these peaks, we focused on the peak on chromosome 11, revealing an insertion of a 6.2-kb *Copia*-type retrotransposon upstream of SIN_1004470, a UDP-glycosyltransferase (*UGT*) gene. In Arabidopsis, overexpression of the *UGT* genes *UGT73C5* or *UGT73C6* has been reported to cause a brassinosteroid-deficient phenotype accompanied by delayed flowering[55, 56], supporting the idea that altered regulation of SIN_1004470 expression may contribute to variation in flowering time in sesame. Sesame accessions carrying this *Copia* insertion near SIN_1004470 flowered significantly earlier than those without the insertion (Fig. 2e, right).

To investigate whether the *Copia* insertion influenced gene expression, we performed RT-qPCR to assess SIN_1004470 expression in an early-flowering accession carrying the insertion (IT216140) and a late-flowering accession without the insertion (IT201452). SIN_1004470 expression levels were constant before and after flowering in IT216140, whereas SIN_1004470 was significantly upregulated after flowering in IT201452 (Fig. 2g), suggesting a *cis*-regulatory influence of the *Copia* insertion located upstream of SIN_1004470.

To experimentally validate the above flowering-associated structural variants (SVs) and develop practical genotyping assays, we designed PCR markers with three primers targeting each SV identified on chromosomes 6 and 11 (Fig. 2d, f, h; Additional file 2: Table S6). The chromosome 6 marker distinguished the two alleles based on amplicon size, producing a 240-bp fragment for the intact allele and a 410-bp fragment for the allele carrying the deletion. Agarose gel electrophoresis showed the expected allele-specific banding patterns, consistent with the SV genotypes inferred from reference genome–based *in silico* prediction analysis (Fig. 2h, left). For the SV on chromosome 11, the marker was expected to generate a 584-bp product for the allele with the *Copia* insertion and a 785-bp product for the allele without the insertion. Indeed, we detected the expected 584-bp band in accessions carrying the *Copia* insertion. In accessions lacking the insertion, however, we detected an approximately 1.2-kb band in addition to the expected 785-bp product (Fig. 2h, right). Because this larger amplicon was approximately 430 bp longer than expected, we performed Sanger sequencing on the 1.2-kb PCR products to determine whether it resulted from non-specific amplification or an additional sequence variation at the target locus. Sanger sequencing identified a 437-bp insertion within the target region between the sites to which the primers com-F and com-R annealed, thus reflecting an additional insertion polymorphism distinct from the *Copia* insertion. Collectively, these results indicate that the PCR-based markers developed in this study can be used for rapid detection of flowering time–associated SVs in sesame accessions.

Notably, for the flowering-associated alleles, both intact forms (i.e. the allele not carrying the deletion on chromosome 6 and the allele not carrying the *Copia* insertion on chromosome 11), were associated with later flowering time. These alleles were enriched in groups G2-1, G2-3, and G3, which are predominantly distributed in regions close to the equator (Figs. 1a,b, 2i). This geographical distribution suggests that low-latitude sesame accessions represent an informative resource for dissecting the genetic basis of flowering time regulation.

### High lignan content in sesame seeds is associated with newly identified genomic variants

Seed lignan contents across the 300 sesame accessions exhibited wide variation, with strong positive correlations among the contents of the three lignan metabolites tested here (Fig. 3a, Additional file 1: Fig. S4b, and Additional file 2: Table S4). We detected genetic variants associated with these traits on chromosomes 6, 11, and 13 (Fig. 3b–d). On chromosome 6, we detected two significant association peaks via *k*-mer GWAS, with the same *k*-mer mapping to the first exons of SIN_1020904 and SIN_1020906, respectively (Fig. 3c,e). This *k*-mer captured the same nonsynonymous substitution in both genes encoding related members of the glycosyl hydrolase (GH) family of enzymes.

**Figure 3.**
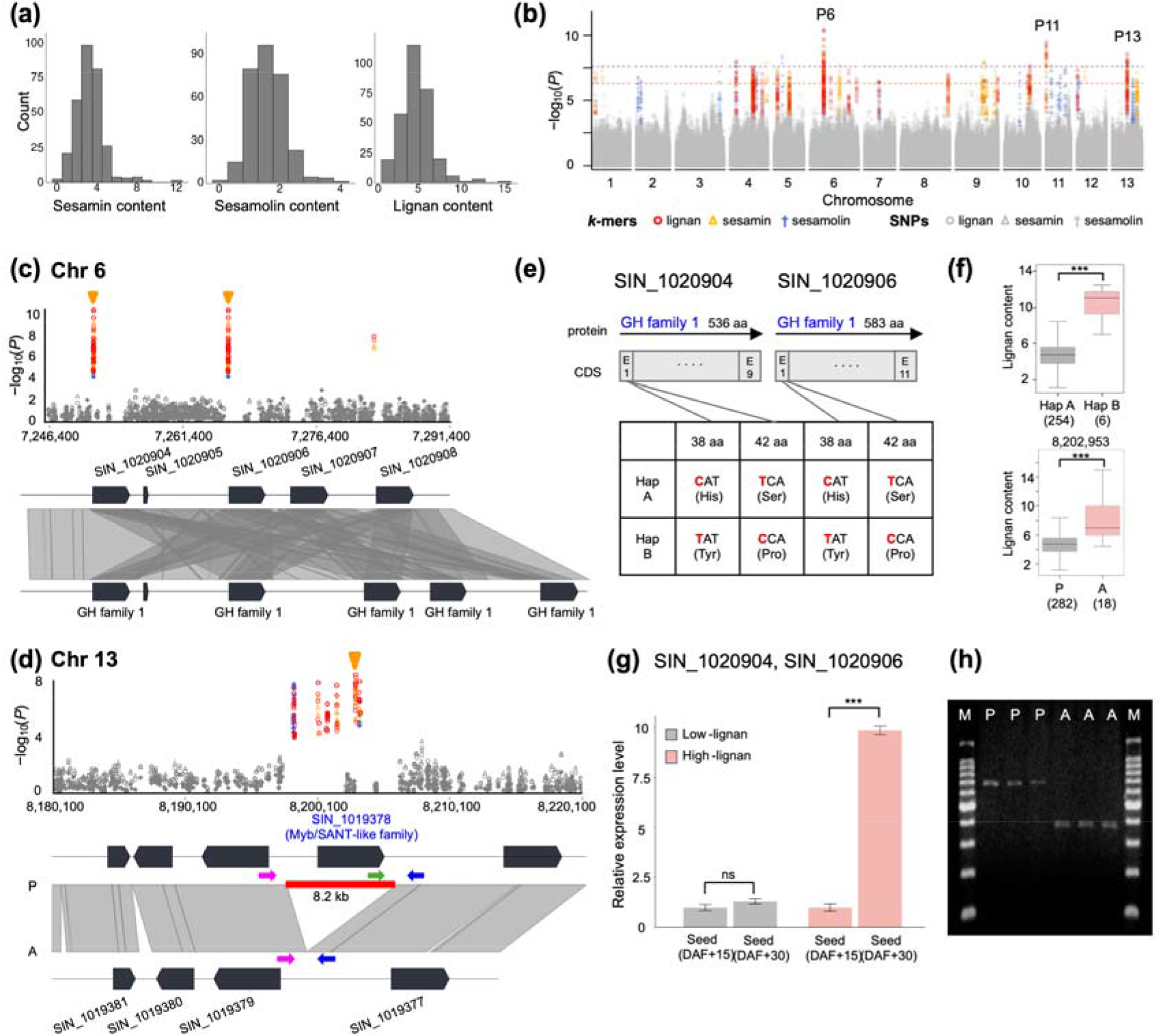
Phenotypic variation and identification of novel single-nucleotide polymorphisms and structural variants associated with lignan content. (a) Variation in sesamin, sesamolin, and total lignan content across two years. (b) Combined Manhattan plot of SNP- and *k*-mer-based GWAS using two-year mean values of sesamin, sesamolin, and total lignan content. Purple and pink dashed lines indicate Bonferroni thresholds for SNPs (*P* = 2.5 × 10**□□**) and *k*-mers (*P* = 5 × 10**□□**), respectively. (c) Two *k*-mer association peaks on chromosome 6 identified the same set of variants, including two nonsynonymous substitutions in both SIN_1020904 and SIN_1020906, and a copy number variant involving nearby glycosyl hydrolase (GH) encoding genes. Orange arrows at Chr6:7,251,389 bp and Chr6:7,266,528 bp indicate the start positions of the most significant *k*-mers within the left and right peaks, respectively. (d) A structural variant (SV) on chromosome 13 was an 8.2-kb deletion encompassing the entire SIN_1019378 locus. The orange arrow at Chr13:8,202,953 bp indicates the start position of the significant *k*-mer. Primer arrows indicate non-deletion-allele-specific forward (green), common forward (pink), and common reverse (blue) primers. (e) Gene structures of SIN_1020904 and SIN_1020906 with nonsynonymous substitutions annotated at nucleotide and amino acid levels. (f) Boxplots of total lignan content by haplotypes A and B defined in (e) (top), and by presence /absence of the significant *k*-mer within the SIN_1019378 region on chromosome 13 (bottom). Two-sided Wilcoxon rank-sum tests; \**P* < 0.05, \*\**P* < 0.01, \*\*\**P* < 0.001. (g) Relative expression of SIN_1020904 and SIN_1020906 in seeds at 15 and 30 days after flowering (DAF) from low-lignan IT318661 (grey) and high-lignan IT169250 (pink). Means ± SE of three biological replicates. Student’s *t*-test; ns, not significant; \**P* < 0.05, \*\**P* < 0.01, \*\*\**P* < 0.001. (h) Gel electrophoresis of markers detecting the SV on chromosome 13 in low- and high-lignan genotypes. A 100-bp DNA ladder was used; expected amplicons are 680 bp (non-deletion) and 374 bp (deletion).

The combination of these two polymorphisms defined two major haplotypes (Fig. 3e). Accessions carrying haplotype B in SIN_1020904 and SIN_1020906, which encodes proteins with Tyr-38–Pro-42, exhibited higher lignan contents than those carrying haplotype A in the same genes, which encode proteins with His-38–Ser-42 (Fig. 3f, top). In addition, the assembled CRR547930 genome showed an approximately 15.9-kb copy number variation (CNV) region encompassing neighboring *GH* genes within this region (Fig. 3c). To evaluate this CNV across the global panel of 300 sesame accessions, we examined sequencing depth for four GH genes (SIN_1020904, SIN_1020906, SIN_1020907, and SIN_1020908) in this region (Additional file 2: Table S7). The mean sequencing depth in six accessions carrying haplotype B was approximately 1.5-fold higher than that in the other 254 accessions carrying haplotype A (Additional file 2: Table S7), consistent with a greater gene copy number in haplotype B. Furthermore, expression analysis showed that these *GH* genes are significantly more highly expressed in mature seeds from the high-lignan accession IT169250 carrying haplotype B compared to the low-lignan accession IT318661 carrying haplotype A (Fig. 3g). These results suggest that higher GH gene copy number and expression may contribute to enhanced lignan accumulation in sesame seeds.

An additional region on chromosome 13 was characterized by an approximately 8.2-kb deletion spanning SIN_1019378, which encodes a MYB/SANT-like transcription factor; the deletion of this gene was associated with higher seed lignan content (Fig. 3d, 3f bottom). MYB transcription factors are central regulators of phenylpropanoid metabolism in plants, and several MYB proteins have been reported to repress the expression of genes within this pathway and modulate metabolic flux[57, 58]. The association between the 8.2-kb deletion across SIN_1019378 and higher lignan content therefore suggests that this MYB-like factor may act as a negative regulator of lignan and phenylpropanoid biosynthesis. To experimentally validate the lignan-associated SV identified on chromosome 13 and develop a practical genotyping assay, we designed a PCR marker targeting this locus (Fig. 3d, h, and Additional file 2: Table S6). We performed PCR amplification using representative high-lignan accessions carrying the deletion and low-lignan accessions carrying the intact locus. The marker was expected to generate a 680-bp amplicon from the intact allele and a 374-bp amplicon from the allele with the deletion. Agarose gel electrophoresis showed the expected allele-specific banding patterns, consistent with the SV genotypes inferred from reference genome–based *in silico* prediction (Fig. 3h).

Within the region on chromosome 11, significant *k*-mer peaks highlighted SIN_1005755 (*NAC SECONDARY WALL THICKENING PROMOTING FACTOR 1* [*SiNST1*])[11, 25] and SIN_1005756[11, 59], both previously implicated in lignan biosynthesis (Fig. 3b). Earlier studies also identified DIRIGENT PROTEIN (DIR) family members, including SiDIR21, together with the sesame-specific PIPERITOL/SESAMIN SYNTHASE (PSS), as candidate components of the lignan biosynthetic pathway[60, 61]. Taken together, these results suggest that high lignan content in sesame is associated with multiple classes of genomic variation, including coding changes, copy number variation, and deletion of a putative regulatory factor.

### Abiotic stress and seed-coat color-related genes are genetically linked to antioxidant capacity in sesame seeds

Sesame seeds are rich in lignans and phenolic acids, and exhibit high antioxidant capacity related to reactive oxygen species scavenging and suppression of lipid oxidation in foods[47, 62, 63]. Given this pharmacological and industrial relevance, we quantified variation in antioxidant capacity across the global collection of 300 sesame accessions using total phenolic content (TPC) and antioxidant activities measured by ABTS (2,2-azino-bis-3-ethylbenzothiazoline-6-sulfonic acid) and DPPH (2,2-diphenyl-1-picrylhydrazyl) assays. All three traits showed wide variation and strong positive pairwise correlations (Fig. 4a, Additional file 1: Fig. S4c, and Additional file 2: Table S4). GWAS based on both SNPs and *k*-mers identified a significant signal on chromosome 6 (Fig. 4b,c). The associated interval between 18,593,000 bp and 18,624,000 bp contains four annotated genes, SIN_1023226, SIN_1023227, SIN_1023228, and SIN_1023229. Within this region, nonsynonymous variants at two sites in SIN_1023226 defined two major haplotypes and nonsynonymous variants at three sites in SIN_1023227 also defined two major haplotypes for that locus (Fig. 4c). Accessions harboring haplotype A (encoding Arg-494–Ser-542) of SIN_1023226, annotated as a WRKY transcription factor gene[64], showed higher antioxidant activity than those with haplotype B encoding Thr-494–Arg-542 (Fig. 4d). Similarly, accessions carrying haplotype A (encoding Ile-47–Thr-580–Thr-667) at SIN_1023227, annotated as a cation/H**O** antiporter gene, displayed higher antioxidant activity than accessions with haplotype B encoding Val-47–Ser-580–Ile-667 (Fig. 4e). Notably, previous studies reported that expression of SIN_1023226, named *SiWRKY67* in the study, responds to waterlogging and drought stress[64, 65]. In addition, SIN_1023227 encodes a predicted cation/H**O** exchanger belonging to a family implicated in pH homeostasis and abiotic stress responses[66, 67]. These observations suggest a potential link between stress-response pathways and antioxidant capacity in sesame. Moreover, SIN_1023226 and SIN_1023227 have been proposed as candidate genes underlying variation in seed-coat color in sesame[68, 69]. Given that flavonoids, phenolic acids, and related secondary metabolites differ among sesame accessions with different seed-coat color and influence antioxidant properties[70–72], variation in seed-coat color may contribute to variation in antioxidant capacity. Taken together, these findings suggest that these antioxidant-associated loci may connect antioxidant capacity with variation in both abiotic stress responses and seed-coat color in sesame.

**Figure 4.**
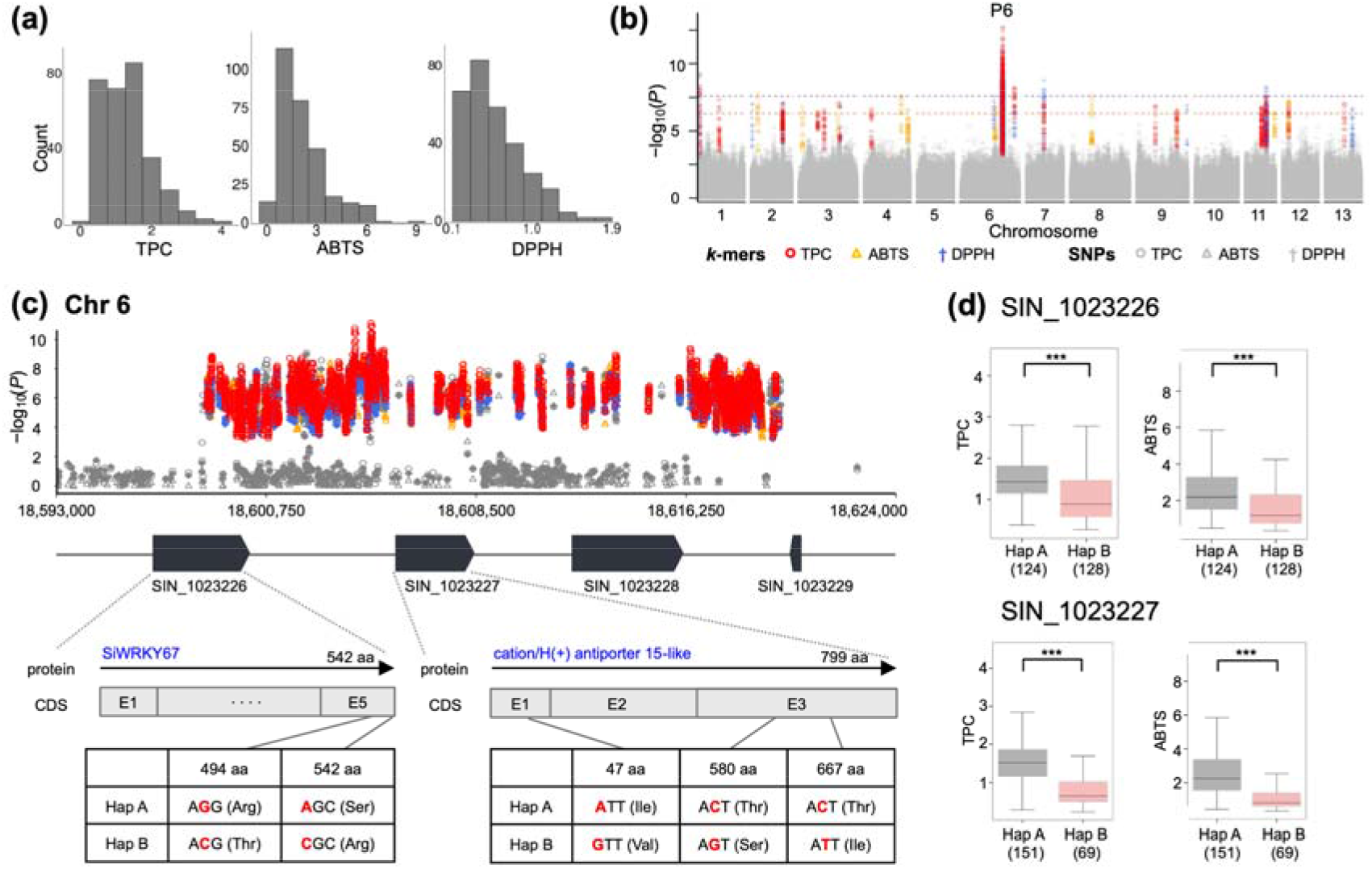
Phenotypic variation and candidate genes associated with antioxidant activities. (a) Phenotypic variation in total phenolic content (TPC), 2,2-azino-bis-3-ethylbenzothiazoline-6-sulphonic acid (ABTS) radical scavenging activity and 2,2-diphenyl-1-picrylhydrazyl (DPPH) radical scavenging activity measured over two years. (b) Combined Manhattan plot of SNP-and *k-*mer-based GWAS for the two-year mean values of TPC, ABTS, and DPPH. The purple and pink dashed lines indicate the Bonferroni thresholds for SNPs (*P* = 2.5 × 10**□□**) and *k*-mers (*P* = 5 × 10**□□**), respectively. (c) Local Manhattan plot of the SNP/*k*-mer association hotspot on chromosome 6, including two candidate genes, SIN_1023226 and SIN_1023227. Gene structure diagrams are shown below the association plot, with nonsynonymous variants used to define gene-specific haplotypes. (d) Haplotype-based comparisons between haplotypes A and B for SIN_1023226 (upper row) and SIN_1023227 (lower row). For each gene, the left and right boxplots show TPC and ABTS radical scavenging activity, respectively. Two-sided Wilcoxon rank-sum tests; \**P* < 0.05, \*\**P* < 0.01, \*\*\**P* < 0.001.

## Discussion

The global sesame panel established in this study provides a valuable genetic resource by revealing previously unreported genomic associations with flowering time, seed lignan content, and antioxidant capacity. These findings broaden the allelic diversity available for improving phenological adaptability and seed quality under increasingly variable seasonal environments.

Among the loci newly uncovered in this study, several variants derived from large InDels or CNVs were associated with flowering time and lignan content; an additional locus carrying nonsynonymous polymorphisms was associated with antioxidant capacity (Fig. 5). These results demonstrate that key trait variation in sesame cannot be fully explained by SNPs alone. Rather, structural variation substantially contributes to allelic diversity at complex genomic regions. Therefore, improving the resolution of these regions across diverse accessions could advance our understanding of the genetic mechanisms underlying metabolic regulation and agronomic trait variation in this crop.

**Figure 5.**
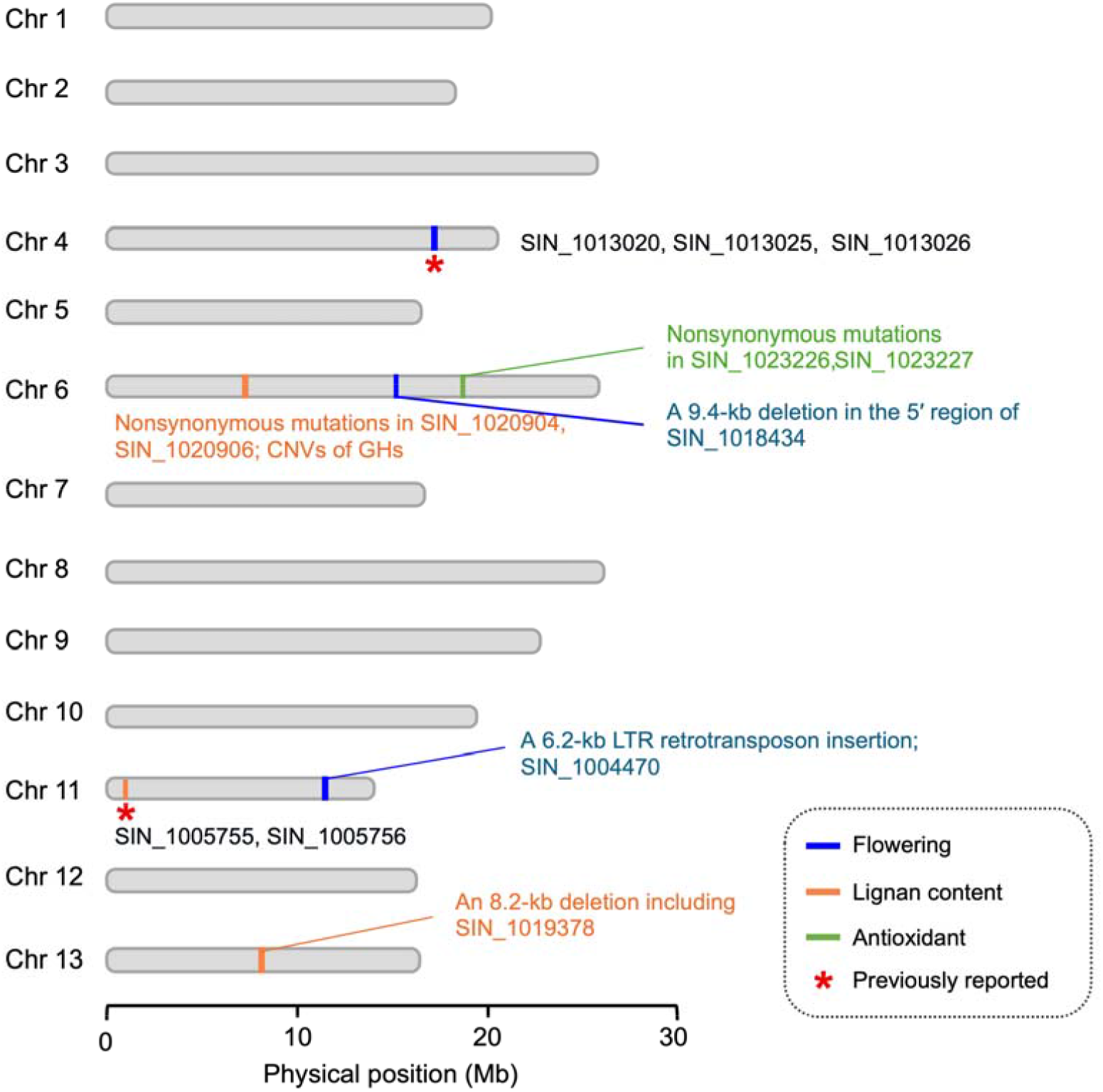
Chromosomal distribution of association loci for flowering-related traits, lignan content, and antioxidant activities in sesame. The loci identified in this study are shown according to trait category: flowering-related traits, blue; lignan content, orange; antioxidant activities, green. Red asterisks indicate loci overlapping previously reported candidate genes.

The flowering time–associated SVs detected here were relatively enriched among accessions originating from lower-latitude regions. This observation suggests that geographically structured allelic distribution may have contributed to adaptive variation in flowering time. In terms of seed-quality traits, the complex CNV associated with lignan content on chromosome 6 was not fully resolved, but it demonstrates how significant variation can be present in genomes that would be difficult to detect or interpret using conventional SNP-based analysis. Such genomic complexity further illustrates why a sesame pangenome is needed, and this globally diverse panel could serve as a useful resource for selecting representative accessions for its construction.

## Conclusions

Taken together, these findings highlight the value of combining globally diverse accessions with SV-aware analysis to uncover alleles associated with variation in flowering time and/or seed-quality traits that may be missed by SNP-based approaches, with implications for breeding plants to thrive in increasingly variable environments.

## Methods

### Plant materials and phenotyping

A natural population of 300 globally representative sesame accessions was selected from the Rural Development Administration (RDA)-Genebank for this study (Additional file 1: Fig. S1 and Additional file 2: Table S1). These accessions were cultivated over three years (2021, 2022, and 2024) at the RDA-Genebank field (35°49**□**53.76**″** N, 127°3**□** 50.4**″** E) using a randomized complete block design. Four agronomic traits were characterized across all three seasons using modified descriptors from the International Plant Genetic Resources Institute (IPGRI)[73]: days to flowering (DTF), days to maturity (DTM), height to the first capsule (HTFC), and capsule zone length (CZL). DTF was recorded as the number of days from sowing until 50% of the plants had initiated flowering (based on the presence of at least one open flower), and DTM as the number of days until 75% of the plants had reached physiological maturity (based on the onset of yellowing of the leaves and stem). HTFC and CZL were measured from five individual plants per accession, as the length (cm) from the base to the first capsule-bearing node, and from the first to the last capsule-bearing node, respectively. Following two harvest seasons (2021 and 2022), seed lignan contents, specifically sesamin and sesamolin, were quantified using high-performance liquid chromatography (HPLC) based on modified protocols[43, 74]. Measurements were performed using three biological replicates per accession, and total lignan content was calculated as the sum of sesamin and sesamolin contents. Antioxidant activity, based on total phenolic content (TPC, µg gallic acid equivalents [GAE]/mg sample), 2,2**′**-azino-bis(3-ethylbenzothiazoline-6-sulfonic acid) (ABTS) radical scavenging activity (µg Trolox equivalents [TE]/mg sample), and diphenyl picrylhydrazyl (DPPH) radical scavenging activity (µg ascorbic acid equivalents [AAE]/mg sample), was also determined in both seasons using an Eon Microplate Spectrophotometer with three biological replicates per accession, following a previously established protocol[44].

### Sampling and sequencing

One representative plant from each accession was chosen for genome sequencing. Genomic DNA was extracted from the young leaves of 14-day-old seedlings using a Gentra Puregene Tissue Kit (QIAGEN, Inc., Valencia, CA, USA), following the manufacturer’s instructions. The sequencing libraries were constructed from high-quality DNA using a TruSeq Nano DNA HT Sample Preparation Kit (Illumina, USA), with an insert size of ∼ 350 bp. Sequencing of each library was performed on the Illumina NovaSeq 6000 platform (Illumina, San Diego, CA, USA), generating paired-end 150-bp reads.

### Variant calling from 300 global sesame germplasm

All raw paired-end reads were pre-processed into high-quality reads (Phred score > 20, > 50 bp) using Trimmomatic (v0.39)[75]. This step involved the removal of adapter sequences and low-quality sequences using the parameter “ILLUMINACLIP:adaptor:2:30:10 SLIDINGWINDOW:4:20 LEADING:3 TRAILING:3 MINLEN:50”. The cleaned reads were then mapped to the ‘Zhongzhi No. 13’ reference genome, version 2[48], using the Burrows-Wheeler Aligner program[76] with default parameters. To remove duplicate PCR reads, the Dedup algorithm of Sentieon was used for deduplication. SNPs and small insertions and deletions (InDels) were called from BAM files using the Sentieon software package (v.202112.01) [77], which is built on the GATK variant calling tool (v4.4.0.0)[78]. Variants for each sample were called using the Sentieon Haplotyper function, followed by joint variant calling with the GVCFtyper function in Sentieon. Variant annotation was conducted using SnpEff (v5.0e)[79]. Raw SNPs and small InDels were filtered with GATK using the following criteria: ‘QD < 2.0 || FS > 60.0 || MQ < 40.0 || MQRankSum < −12.5 || ReadPosRankSum < −8.0’ for SNPs and ‘QD < 2.0 || FS > 200.0 || ReadPosRankSum < −20.0’ for small InDels. Filtered biallelic variants were retained if they were supported by 5 to 100 reads, had missing genotype rates below 20%, and had a minor allele frequency above 5%. The resulting VCF file was used for subsequent analysis.

### Population structure, phylogenetic, and principal component analyses

A total of 1,965,411 SNPs located across the chromosomes of the sesame genome were retained for population structure, phylogenetic, and principal component analyses. These SNPs met the criteria of a minor allele frequency above 5% and a missing genotype rate below 20%. Population structure analysis was conducted using ADMIXTURE (v1.3.0)[80], exploring cluster values (*k*) ranging from 2 to 20 and performing ten runs with random seeds for each *k*. The ADMIXTURE results were visualized using Pophelper[81]. Phylogenetic analysis was performed using FastTree (v2.1.11)[82] with the general time reversible model, and the resulting tree was visualized using FigTree (v1.4.4)[83]. Principal component analysis (PCA) was performed using PLINK v1.90b6.21[84].

Additionally, to evaluate the diversity of the 300 sesame accessions, all 1,368 available whole-genome sequencing datasets consisting of paired-end Illumina reads for sesame (Additional file 2: Table S8) were downloaded from the NCBI Sequence Read Archive (SRA)[85] using fastq-dump tool from the SRA Toolkit[86]. These SRA datasets were selected by searching for “*Sesamum indicum*” and applying filters for “Source–DNA, Library Layout–Paired, Platform–Illumina, and Strategy–Genome Sequencing” on the NCBI SRA website[85]. PCA was then performed on the combined dataset comprising 1,668 accessions, consisting of SNP data from the 300 sesame accessions sequenced in this study and the 1,368 SRA datasets, using 1,698,272 SNPs.

### Regional genetic overlap

To compare gene pools between regions rather than between individuals, genetic overlap was quantified with ReMIXTURE v0.1.0 in R v4.4.2. The VCF file generated as described above was converted into a pairwise allele-sharing distance matrix (1 − IBS) in PLINK v1.90b6.21[84]. Individuals were labeled by their region of origin, and ReMIXTURE was run for 1,000 iterations over the series of smoothing bandwidths (H) selected automatically by the package (random seed 42).

### Linkage disequilibrium decay analysis

Linkage disequilibrium (LD) was analyzed in the 300 sesame accessions by calculating pairwise *r^2^*, the squared correlation coefficient between allele frequencies at two loci, for 1,965,411 SNPs distributed across the 13 chromosomes of the sesame genome using PopLDdecay software[87]. LD values were estimated for the entire population and for each chromosome. The physical distance at which *r^2^* dropped to half of its maximum value was determined and used to define LD decay.

### Genome-wide association study

A total of 2,032,196 high-quality SNPs, encompassing all variants across chromosomes and scaffolds, were used for SNP-based GWAS, which was performed using the GAPIT program[88] with the mixed linear model. For the *k*-mer-based GWAS, Illumina short-read sequencing data from the 300 sesame accessions were fragmented into 31-mer segments, and a presence/absence table of 31-mers was constructed following the pipeline by Voichek and Weigel[89]. GWAS was conducted using 100,000 *k*-mers with the kmers_gwas.py script[89] and GEMMA (v0.98) software[90], applying a minor allele frequency threshold of 5%. Selected *k*-mers were aligned to the reference sesame genome[48] using BLASTN, retaining only complete 31-bp alignments. Among the selected *k*-mers, those with significant −log_10_ *P*-values exceeding 6.3 were chosen for clear signal identification. Three field trials were conducted in 2021, 2022, and 2024 to characterize flowering-related traits. The trial data were analyzed using the lme4 package[91] in R to generate best linear unbiased estimates (BLUEs), which were subsequently used as input for GWAS. For lignan content and antioxidant activities, the average evaluation data from two years (2021 and 2022) were used as input for GWAS.

### Genome assembly and structural variation analysis

Genome assembly was conducted using previously published long-read sequencing data for sesame[41], including four Nanopore datasets (CRR547927, CRR547928, CRR547929, CRR547930) and one PacBio dataset (CRR565260), obtained from the Genome Sequence Archive at the China National Center for Bioinformation[92]. *De novo* genome assemblies for the five sesame accessions were generated with Flye using default parameters[93]. A graph-based genome was constructed from these five datasets using Minigraph-Cactus[94]. Candidate regions with structural variants corresponding to the *k*-mer-specific GWAS signals were inspected and validated using the odgi toolkit[95].

### Expression analysis

To examine gene expression differences associated with flowering time, upper leaves were sampled from an early-flowering genotype (IT216140, 43.7 days) and a late-flowering genotype (IT201452, 100.3 days) at two developmental stages: before flowering and immediately after flowering. To investigate differences in expression associated with lignan content, seeds within capsules were collected from a low-lignan genotype (IT318661, 1.21 mg g**□** ¹) and a high-lignan genotype (IT169250, 12.40 mg g**□** ¹) at 15 and 30 days after flowering. Total RNA was extracted from each sample using TRIzol reagent (Invitrogen, CA, USA) according to the manufacturer’s instructions. The RNA was reverse-transcribed into cDNA using a ReverTra Ace qPCR RT Master Mix with gDNA Remover kit (Toyobo, Japan) following the manufacturer’s protocol. Primers for RT-qPCR were designed using Primer3[96] (Additional file 2: Table S9). Quantitative PCR (qPCR) was performed with gene-specific primers using TOPreal™ qPCR 2X PreMIX (Enzynomics, Korea) on a LightCycler® 96 System (Roche, Germany). The amplification conditions consisted of an initial denaturation at 95°C for 10 min, followed by 50 cycles of 95°C for 10 s, 58°C for 10 s, and 72°C for 10 s. Each assay was conducted in triplicate using independent biological samples. The sesame *Actin7* gene (SIN_1006268) was used as the internal control. Relative expression levels were calculated using the 2^−ΔΔCT^ method. Statistical significance of differences between groups was assessed using Student’s t-test.

## Declarations

### Ethics approval and consent to participate

Not applicable

### Consent for publication

Not applicable

### Availability of data and materials

Whole-genome resequencing data generated from 300 sesame accessions during the current study are available in the NCBI Sequence Read Archive (SRA) under BioProject accession PRJNA1311236. The code used for the *k*-mer-based GWAS workflow is available on GitHub (https://github.com/whyskyisgray/Sesame_kmer-based_GWAS).

### Authors’ contributions

S.L. designed the study. S.L., J.-E.L., and S.K.L. performed phenotyping. S.L. and S.H.L. performed the GWAS analyses. J.-Y.A. performed the RT-qPCR experiments. S.L., S.H.L., and J.-Y.A. analyzed the resulting data. S.L. and S.H.L. wrote the manuscript. M.J., G.-A.L., and T.-J.Y. reviewed and revised the manuscript. G.-A.L. and T.-J.Y. supervised the project. All authors read and approved the final manuscript.

## Acknowledgements

Not applicable

## Supplementary Information

### Additional file 1: Supplementary figures (Figures S1-S5)

**Figure S1**: Geographic distribution of RDA-Genebank sesame germplasm and selected accessions. (a) Comparison of the geographic distribution between 8,218 accessions from the RDA-Genebank collection and 300 accessions used in this study. (b) Geographic distribution of the 300 selected sesame accessions. ARG, Argentina; BGR, Bulgaria; CHN, China; COL, Colombia; ECU, Ecuador; EGY, Egypt; ETH, Ethiopia; GRC, Greece; IND, India; IRN, Iran; ITA, Italy; JPN, Japan; KEN, Kenya; KOR, South Korea; MEX, Mexico; MOZ, Mozambique; NGA, Nigeria; NIC, Nicaragua; PER, Peru; PHL, Philippines; PRY, Paraguay; RUS, Russia; SDN, Sudan; SEN, Senegal; SOM, Somalia; THA, Thailand; TUR, Türkiye; TWN, Taiwan; USA, United States; UZB, Uzbekistan; VEN, Venezuela. **Figure S2**: SNP density plot across 13 chromosomes in 300 sesame germplasm. The x-axis represents chromosome length in 10-Mb units and the y-axis indicates SNP counts per 10-kb window. SNP density was calculated with a 10-kb window size and a 2-kb step size. **Figure S3**: Principal component analysis of sesame accessions based on PC1 and PC2. Left, principal component analysis (PCA) of the 300 sesame accessions from around the world. Right, PCA of the combined dataset comprising the 300 accessions and 1,368 publicly available accessions from the NCBI Sequence Read Archive (SRA). **Figure S4**: Phenotypic correlations within flowering-related traits, lignan content, and antioxidant activities. (a) Correlations among days to flowering (DTF), days to maturity (DTM), height to the first capsule (HTFC) and capsule zone length (CZL). (b) Correlations among sesamin, sesamolin, and total lignan content. (c) Correlations among total phenolic content (TPC), 2,2-azino-bis-3-ethylbenzothiazoline-6-sulphonic acid (ABTS) radical scavenging activity and 2,2-diphenyl-1-picrylhydrazyl (DPPH) radical scavenging activity. **Figure S5**: Additional variants associated with flowering-related traits identified by GWAS. (a) Manhattan plot showing a SNP/*k*-mer hotspot on chromosome 4 that encompasses known genes and additional candidate genes. Identical significant *k*-mer peaks corresponding to *Copia*-type retrotransposon insertions on chromosome 3 (b) and chromosome 13 (c).

### Additional file 2: Supplementary tables (Tables S1-S9)

**Table S1**: The 300 sesame accessions used in this study. **Table S2**: Summary of resequencing data for the 300 sesame accessions. **Table S3**: Linkage disequilibrium decay distances for individual chromosomes and the whole genome in the 300 sesame accessions. **Table S4:** Summary of the phenotypic dataset used for genome-wide association studies. **Table S5**: Significant SNPs associated with flowering-related traits on chromosome 4 (17,190,000–17,420,000 bp). **Table S6**: Primer sets for detecting structural variants associated with days to flowering and lignan-related traits. **Table S7**: Sequencing depth of GH genes, SNP-based haplotypes, and total lignan content across sesame accessions. **Table S8**: List of 1,368 publicly available sesame sequencing datasets downloaded from the NCBI Sequence Read Archive. **Table S9**: Primers used for RT-qPCR.

